# Augmenting TCR signal strength and ICOS costimulation results in metabolically fit and therapeutically potent human CAR Th17 cell therapy

**DOI:** 10.1101/2022.10.28.514057

**Authors:** Megan M. Wyatt, Logan W. Huff, Michelle H. Nelson, Lillian R. Neal, Andrew R. Medvec, Guillermo O. Rangel Rivera, Aubrey S. Smith, Amalia M. Rivera Reyes, Hannah M. Knochelmann, James L. Riley, Gregory B. Lesinski, Chrystal M. Paulos

## Abstract

Adoptive cell transfer (ACT) therapy with IL-17 producing human T cells elicits potent antitumor activity in preclinical models. However, further refinement of this novel approach is needed to position it for clinical application. While activation signal strength differentially regulates IL-17 production by human CD4^+^ T cells, the degree to which TCR and co-stimulation signal strength impacts antitumor Th17 cell immunity remains poorly understood. We discovered that decreasing TCR/co-stimulation signal strength by incremental reduction of αCD3/co-stimulation beads in a Th17 culture progressively diminished their effector memory phenotype but enhanced their polyfunctionality. Additional investigation revealed that Th17 cells stimulated with αCD3/ICOS beads produced more IL-17A, IFNγ, IL-2 and IL-22 than those stimulated with αCD3/CD28 beads, regardless of signal strength. Th17 cells propagated with 30-fold fewer αCD3/ICOS beads (weak signal strength, 1 bead per 10 T cells) were less reliant on glucose for growth compared to those stimulated with the standard, strong signal strength (3 beads per T cell). Further metabolomic analysis revealed Th17 cells weakly simulated with αCD3/ICOS beads favored the central carbon pathway through increased gluconeogenesis for bioenergetics, marked by abundant intracellular phosphoenoylpyruvate (PEP). Importantly, Th17 cells weakly stimulated with αCD3/ICOS beads and redirected with a chimeric antigen receptor (CAR) that recognizes mesothelin were more effective at clearing large human mesothelioma tumors when infused into mice than those manufactured using the standard FDA-approved protocols. Taken together, these data indicate Th17 ACT therapy can be improved by using fewer activation beads during T cell manufacturing, a finding that is both cost effective and directly translatable to patients.

## Introduction

T cells genetically redirected to express a chimeric or T cell antigen receptor (CAR, TCR) can elicit potent efficacy in a subset of patients with tumors harboring the antigenic target (*1-3*). While CD19-CAR T cell products have demonstrated success for individuals with B cell malignancies, CAR-based therapy approaches have been less effective at treating patients with solid tumors (*4*). Reasons why adoptive T cell transfer (ACT) therapy fails for patients with solid tumors is multi-factorial, including inefficient trafficking of T cells into the tumor (*5*), inability to overcome the oppressive tumor microenvironment (*6*), and low persistence and response maintenance against aggressive malignancies (*4, 7*).

One potential strategy to overcome limitations of ACT therapy is to improve the quality of T cells generated *ex vivo*. Currently, standard FDA-approved ACT protocols recommend logarithmic growth of CAR T cells, which involves the selection of CD3+ T cells from the patient’s blood and expansion with artificial antigen presenting cells (aAPCs). These aAPCs often consist of a magnetic bead coated with αCD3 and αCD28 antibodies to mediate TCR stimulation and co-stimulation, respectively. Three αCD3/CD28 beads are used per T cell in the culture to ensure logarithmic activation and propagation (*8, 9*). As part of these protocols, T cells are genetically engineered to encode an antigen receptor and expanded in the presence of IL-2 for two weeks before reinfusion into the lymphodepleted patient.

We posited that cell therapy against solid tumors can be advanced by generating human Th17 cell products in a less differentiated state. Th17 cells are a helper CD4^+^ T cell subset originally reported as pro-inflammatory and important in mediating defense against self-tissue and infections, via their production of IL-17 (*10, 11*). We and others reported that Th17 subsets elicit potent efficacy against solid tumors in aggressive, pre-clinical tumor models, compared to other helper subsets, including Th1 or Th2 cells (*12-16*). Subsequent investigations revealed Th17 cells possess hallmarks of stemness, marked by enhanced self-renewal, function, and proliferative potential, which make them particularly attractive as a novel ACT approach (*14, 17, 18*). Yet, the ideal way to expand antitumor Th17 cells for ACT without losing these characteristics remains elusive and a significant barrier to their implementation beyond the pre-clinical setting.

Prior work has demonstrated that lowering activation signal strength generates T cells with a younger memory phenotype and higher functionality (*19-21*). We hypothesized that reducing the initial human Th17 activation strength by using fewer αCD3 beads coated with either CD28 or ICOS in the culture would improve their antitumor activity. As signal strength was weakened, fewer Th17 cells differentiated into a full effector memory phenotype. Weak ICOS co-stimulated Th17 cells maintained a more naïve memory phenotype and co-secreted multiple cytokines. Th17 cells stimulated with the standard, strong signal strength relied showed enhanced mitochondrial bioenergetics more on glycolysis for growth and energy, while those stimulated with weak ICOS signal strength had enhanced mitochondrial bioenergetics. Moreover, global metabolomic analysis of these cells identified unique metabolites associated with the central carbon pathway and gluconeogenesis are elevated in Th17 cells generated with weak signal strength ICOS bead preparations. *In vivo*, weak ICOS co-stimulated Th17 cells redirected with a mesothelin-specific CAR persisted and effectively controlled growth of mesothelioma tumors. Our work reveals that modulating metabolic pathways in Th17 cells is possible by simply using fewer beads in a culture, which results in potentiated CAR T cell immunity against solid tumors. These data enable further refinement of *ex vivo* expansion protocols for human Th17 cells that will accelerate the path to future clinical application.

## Results

### Lowering signal strength improves human Th17 cell polyfunctionality

Th17 subsets are more potent against solid tumors compared to other helper subsets, including Th1 or Th2 cells(*13, 14, 22*). We theorized that expanding human Th17 cells with fewer beads would generate lymphocytes with a less exhausted phenotype, marked by their capacity to secrete multiple cytokines at once. To test our hypothesis that reducing TCR/co-stimulation activation signal strength augments Th17 cell functionality and memory phenotype, we serially diluted the number of activation beads added to human Th17 cells to as low as 30-fold less than that used in the clinic, to mimic reduced antigen signaling. Specifically, CD4^+^ T cells were freshly isolated from PBMCs of de-identified healthy human donors, cytokine-polarized toward a Th17-phenotype while being activated with αCD3 magnetic beads coated with either αCD28 or αICOS agonists, as depicted in Figure 1A. Th17-polarized cells were expanded in the presence of IL-2 (100 IU/ml, added 2 days post-polarization) for 10 days and then assayed these end product cells for their capacity to secrete cytokines after PMA and Ionomycin re-stimulation. We found that as signal strength decreased, IFNγ and IL-17A production increased gradually (Figure 1B). Th17 cells expanded with the fewest beads (1:10 bead : T cell ratio) had the greatest percentage of cells that expressed both cytokines, and we observed that this polyfunctionality was most pronounced in cells activated with αCD3/ICOS beads. Note that Th17 cell numbers in cultures were held constant (∼0.8 million/ml in a 24-well plate) while the αCD3/costimulatory bead numbers were altered as a rheostat for activation strength. Our data reveals that decreasing the activator beads profoundly bolsters Th17 polyfunctionality.

**Figure 1:**
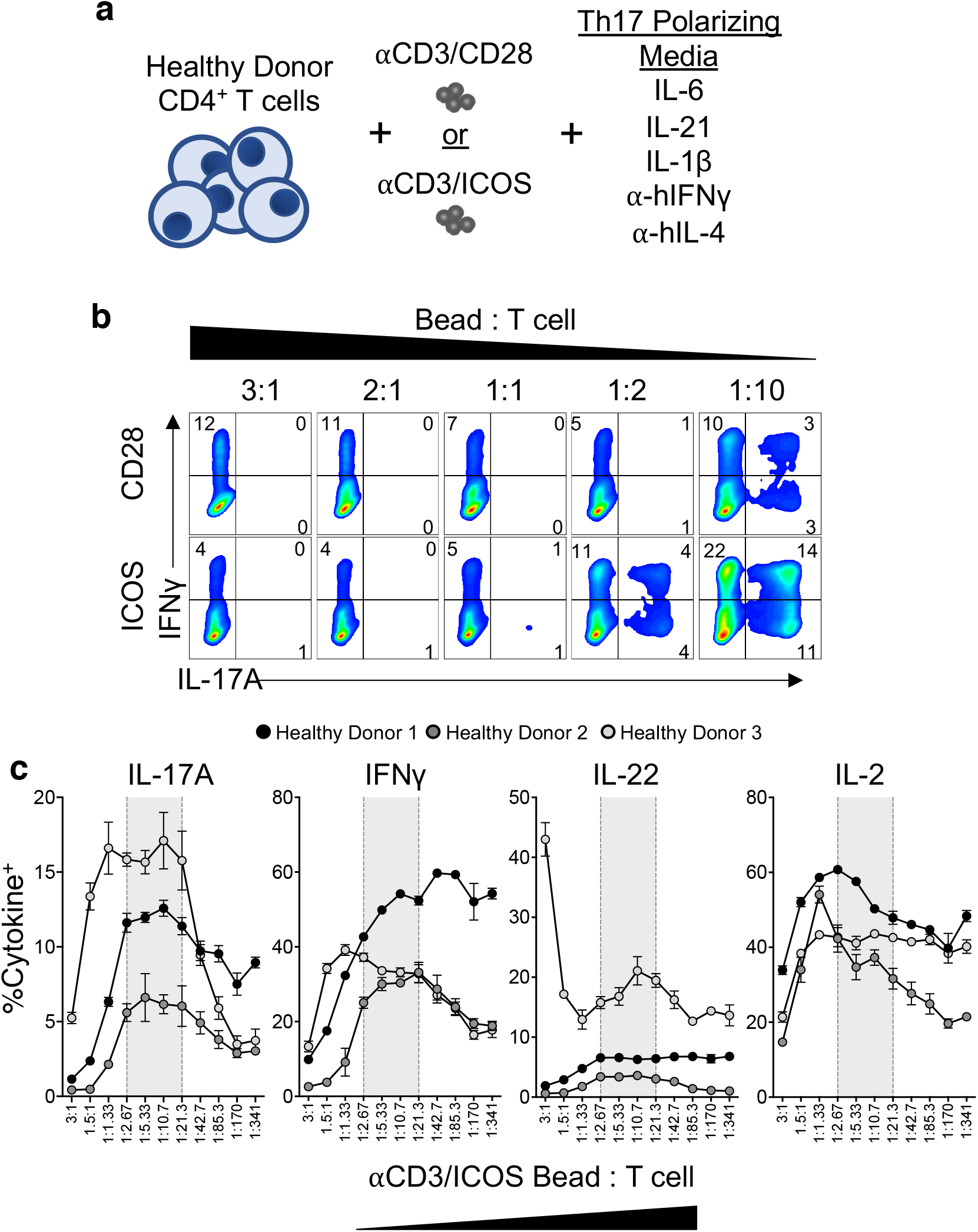
Decreasing activation signal strength results in increased functionality of Th17 cells. a) CD4^+^ T cells from normal human donors were isolated and expanded with either CD3/CD28 or CD3/ICOS artificial APC beads and Th17 polarizing cytokines. b) Percentages of IFNγ and IL-17A cytokine producing Th17 cells expanded for 9 days after activation with either CD3/CD28 (top) or CD3/ICOS (bottom) beads at indicated bead:T cell ratios, as assessed by flow cytometry. c) Percentages of IL-17A, IFNγ, IL-22 and IL-2 cytokine producing Th17 cells activated with a range of T cell : αCD3/αICOS bead ratios as determined by flow cytometry. Shaded area indicates approximate range of observed optimal cytokine production (n=3 normal donors, 3-8 technical replicates per donor)

To uncover the ideal number of beads needed to optimize IL-17, IFNγ, IL-22 and IL-2 production by ICOS co-stimulated Th17 cultures, cells were cultured in a more dynamic titration range of activator beads (αCD3/ICOS) and then assessed for their functional profile (Figure 1C). Cytokine production peaked in T cells stimulated at an intermediate T cell to bead ratio (∼1:3 – 1:21 bead per T cell). Conversely, Th17 cells expanded via strong signal strength using the standard ratio (3:1 bead per T cell) produced significantly lower levels of all cytokines tested. At lower signal strengths of ∼1:43 bead: T cell and below, cytokine production gradually tapered, albeit their production of any 4 cytokines remained higher than those expanded with high signal strength cells (i.e., ∼3:1 – 1:1 bead per T cell). Overall, our data reveal an optimal range in which fewer beads can be used to manufacture Th17 cell products with enhanced multifunctionality.

### Lowering activation signal strength slows Th17 cell division but sustains naïve memory phenotypes

Current ACT protocols require expansion of large numbers of T cells before they are infused back into patients. Proliferation assays were done to test if reducing the number of beads added to Th17 cells would impact their expansion and yield by culturing them with either a “Weak” (1 bead per 10 T cells) or “Strong” bead stimulus protocol (the standard 3 beads per T cell). In Th17 cultures activated with αCD3/CD28 beads, we found that both the Strong and Weak bead per T cell groups divided within 2 days (Figure 2A). By day 4, all cells in the CD28 Strong and CD28 Weak groups divided at least once. One week later, most cells in both groups had divided. In contrast, αCD3/ICOS stimulated Th17 cells lagged in proliferation compared to those stimulated with αCD3/CD28 beads at the same ratios. In fact, by day 7 post-activation, a small percentage of ICOS Weak cells did not divide. These data suggest a difference in ICOS and CD28 signaling cues in promoting T cell proliferation, potentially due to the constitutive presence of CD28 on the T cell surface while ICOS upregulates after TCR stimulation (*23*). However, the expansion rates and ultimate cell yield was not significantly different between Th17 cells undergoing either CD28 or ICOS co-stimulation, nor between the ICOS Strong vs. ICOS Weak conditions (Figure 2B). The CD28 Weak condition did significantly lower the yield of Th17 cells by day 10 post-activation, when compared to the CD28 Strong condition (p = 0.0035).

**Figure 2.**
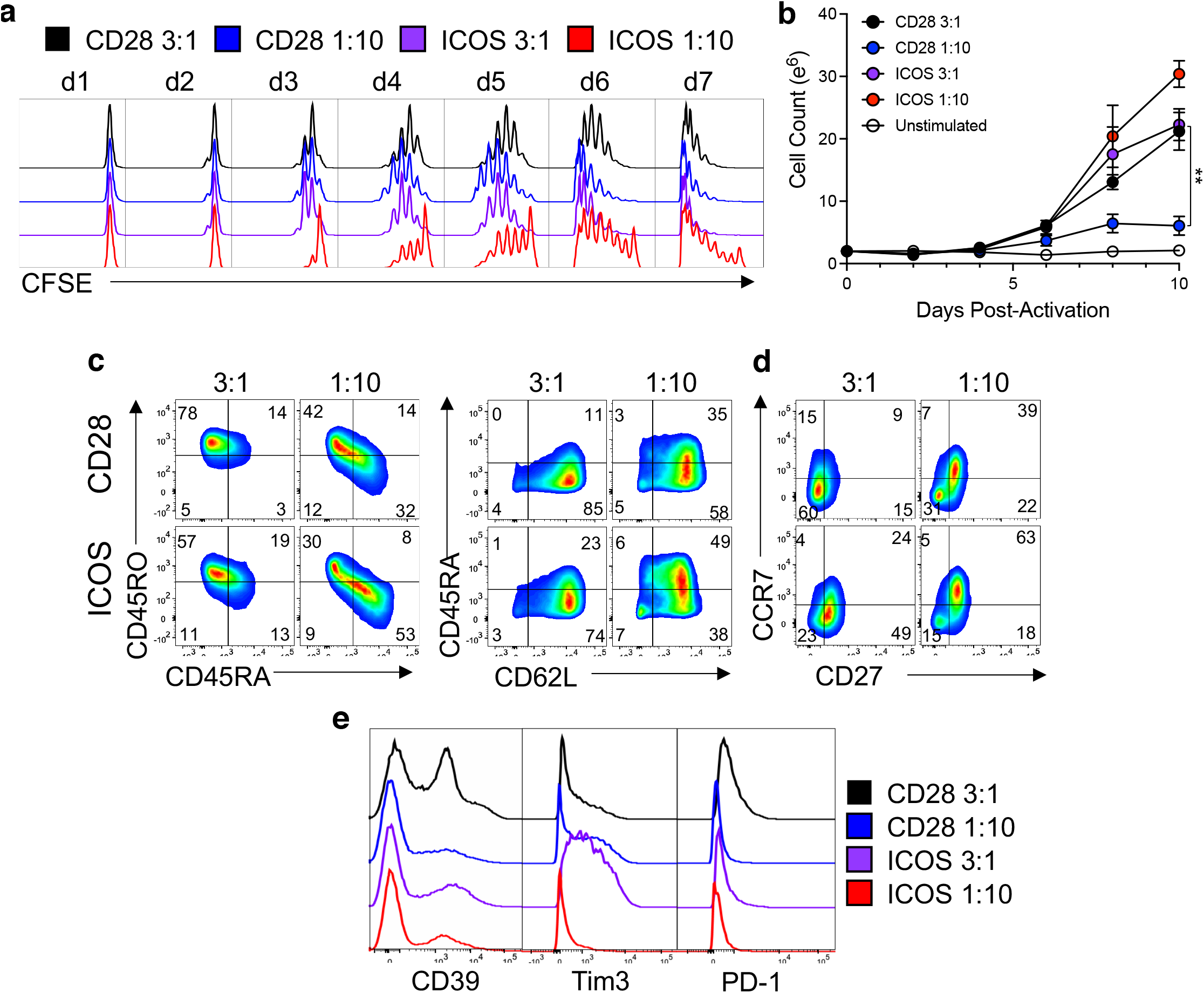
Lower activating signal strength programs an optimal T cell memory phenotype. a) Cell division over one week of culture. CFSE labeled cells were activated with either CD3/CD28 (top) or CD3/ICOS (bottom) beads at three different ratios: 3:1 bead:T cell (black), 1:10 bead:Tcell (blue) or 1:100 bead:T cell (red). Cells from each culture were collected and fixed each day after activation, up to 7 days. After 7 days, all cells were analyzed via flow cytometry. Cell growth rate was also observed (b). Cells from each culture were counted every other day, up to day 10 of culture. Results are graphed as fold change in the number of cells (log_2_). After 10 days of growth, expression of memory markers and co-inhibitory receptors were analyzed by flow cytometry (c). Numbers in the FACS plots represent percent positive of lymphocytes. Statistical analysis was performed using one-way ANOVA with Tukey’s multiple comparisons test and Log-rank Mantel-Cox Test (** =P<0.01).

T cell differentiation and senescence are factors that limit the longevity of T cells used for ACT and hence its success. Thus, it is often a goal of novel manufacturing protocols to produce less differentiated T cell products for infusion. Because T cell expansion drives differentiation, we next assessed the memory profile of Th17 cells expanded with these different stimulation protocols, hypothesizing that higher signal strength (i.e. more beads to T cells) would foster greater effector memory differentiation. As expected, Strong stimulatory conditions promoted the generation of differentiated cells, while Weak signal strength supported more naïve-like memory cells, as indicated by the percent of CD45RA^+^CD45RO^-^ and CD45RA^+^CD62L^+^ cells (Figure 2C). CCR7 and CD27, additional markers expressed on naïve T cells, were also most elevated in Th17 cells given fewer beads (Figure 2D). Expression of T cell exhaustion and coinhibitory markers CD39, Tim3 and PD-1 were all elevated on Strong-stimulated cells (Figure 2E). Thus, we found that weaker signal strength generated Th17 cells were ultimately those with a less differentiated memory phenotype.

### Low signal strength improves Th17 mitochondrial function and reduces utilization of glucose

Mitochondrial health and metabolism are essential to T cell survival, function, and immunity (*24-27*). Consequentially, generating a metabolically fit T cell product is critical for durable antitumor immunity, as it enables resistance to the harsh and nutritionally restrictive tumor environment. Proliferation and T cell differentiation are tightly tied to the commitment to glycolysis (*28*), so we hypothesized that weaker signal strength promoted less glucose reliance in Th17 cells. We assayed the expression of the glucose transporter 1 (GLUT1) and cellular glucose uptake (2-NBDG, a fluorescent glucose analogue) in Th17 cells expanded with titrated αCD3-magnetic beads decorated with CD28 or ICOS agonist antibodies at Strong (3 bead per T cell) and Weak (1 bead per 10 T cells) levels of signal strength over the course of *in vitro* expansion. GLUT1 was consistently elevated in the Strong CD28 activated Th17 cells, with a spike in expression on day 5 of culture (Figure 3A). All other activation treatments expressed less surface GLUT1 throughout culture as compared to the Strong CD28 activated Th17 cells. Concomitantly, Th17 cells were measured for glucose using 2-NBDG. Cells stimulated with a Weak signal strength consumed glucose at a slower rate than those cells activated at the Strong signal strength, regardless of ICOS or CD28 co-stimulatory signaling (Figure 3B). Th17 cells stimulated with the Strong concentration of αCD3/ICOS beads consumed less 2-NBDG than cells expanded with the Strong αCD3/CD28 beads after a week of expansion, suggesting that ICOS co-stimulation decreases T cell reliance on glucose. Due to the observed differences in glucose reliance between the groups, we next probed the relative mitochondrial profile of Weak vs. Strong activated Th17 cells.

**Figure 3:**
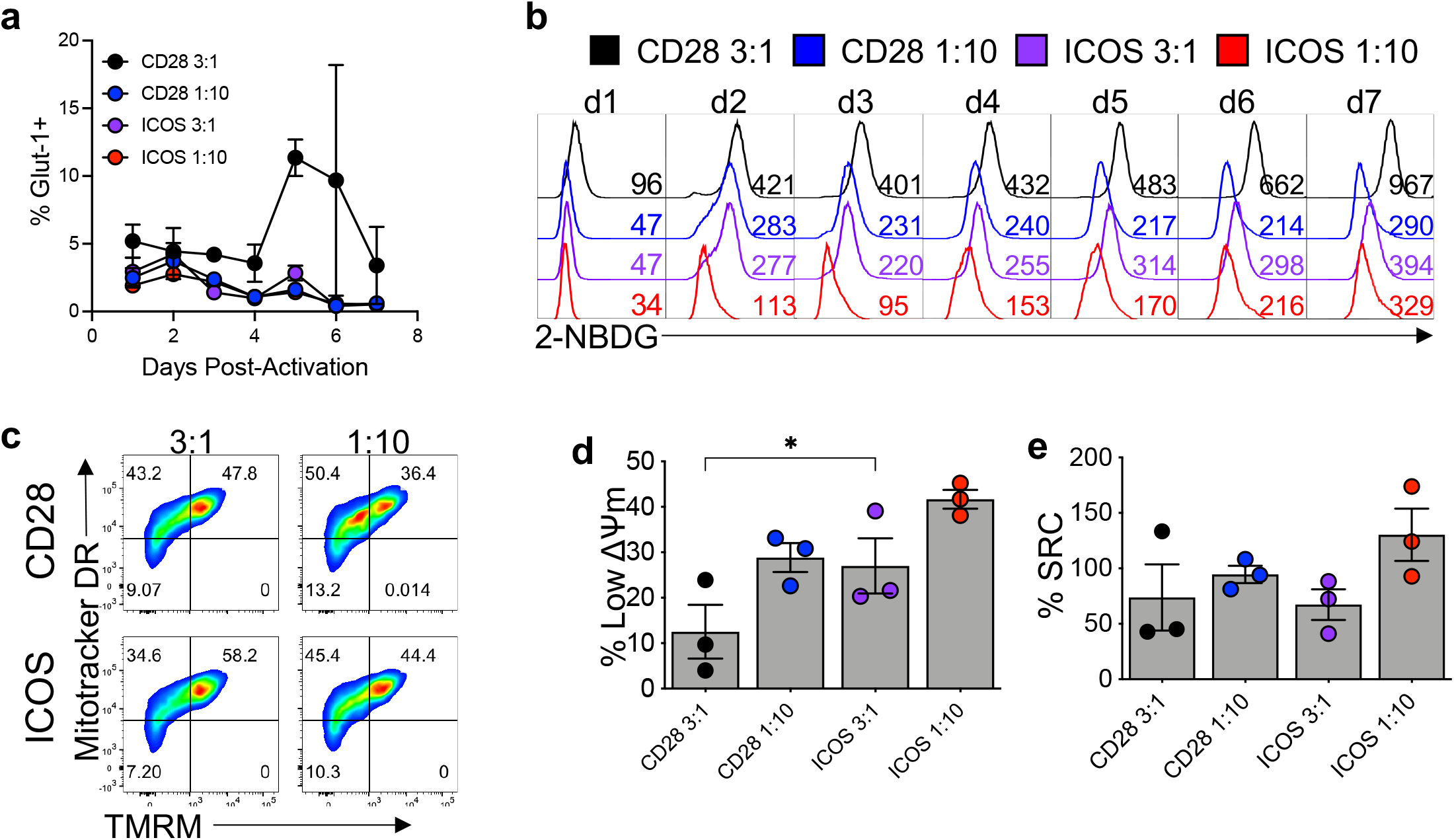
Glucose uptake and mitochondrial dynamics are affected by signal strength and co-stimulatory signal. Flow cytometry was used to evaluate the presence of the GLUT1 receptor on the cells (a) as well as to assess their ability to uptake glucose using the fluorescent glucose reporter 2-NDGB over 7 days of culture (b). At 10 days post-activation, markers of mitochondrial mass (Mitotracker Deep Red) and membrane potential (TMRM) were used to characterize the cells by flow cytometry (c). The percentage of cells with low mitochondrial membrane potential is represented (d). At the same timepoint, a Seahorse mitochondria stress test was used to calculate percent spare respiratory capacity (SRC) (e). For each assay, N=3 healthy donors. Statistical analysis was performed using one-way ANOVA with Tukey’s multiple comparisons test (* = P<0.05 ; ** * =P<0.001).

T cells possessing mitochondria with low mitochondrial membrane potential (ΔΨm) retain enriched markers of stemness and are capable of enhanced tumor ablation *in vivo* (*29*). We hypothesized that weakening signal strength would increase the abundance of Th17 cells with a low ΔΨm signature, based on our findings regarding their memory and polyfunctionality. Th17 cells were profiled for their relative abundance of mitochondria, their mitochondrial phenotype and metabolic capacity. Mitochondrial mass (Mitotracker Deep Red, ThermoFisher) and ΔΨm (TMRM) were analyzed by flow cytometry. As predicted, whether co-stimulated by CD28 or ICOS agonists, lowering the activation signal strength expanded Th17 cells with more of their mitochondria mass exhibiting lower ΔΨm (Figure 3C-D), potentially indicating that Weak signal strength preserves high mitochondrial function and mass. We theorized that the capacity for mitochondrial energy generation in Th17 cells would retain more spare respiratory capacity (SRC) when stressed following cell expansion with fewer activator beads. Using a Seahorse XF assay (Agilent), the percent SRC increased when the activation signal strength was decreased, though not significantly (Figure 3E). Our data imply that mitochondria in Th17 cells activated with a weaker TCR signal strength and receiving ICOS co-stimulatory signaling are less differentiated, proliferate at lower rates and are less reliant on glycolysis *in vitro*. Therefore, Th17 cells stimulated with 30-fold fewer activation beads have altered metabolic requirements than those expanded with more beads commonly used to generate clinical CAR products. Because of the marked decrease in glucose uptake, ability to proliferate and subtle changes in the mitochondrial function, we hypothesized that these Th17 cultures expanded with a reduced signal strength and ICOS co-stimulatory signaling may be altering their metabolic needs to promote their survival and function, so we further probed the metabolic profile of these cells.

### Gluconeogenesis is used by weak TCR activated/ICOS co-stimulated Th17 cells

To interrogate the metabolic profile of human Th17 cells activated with a Strong versus a Weak TCR signal, human CD4^+^ T cells from 7 healthy PBMC donors were programmed towards a Th17 phenotype and then activated, or not, with a Strong (3 beads per T cell) or Weak (1 bead per 10 T cells) number of αCD3/ICOS beads for 4 days, in order to optimally capture the differences between treatments based on our metabolic phenotype time course data (Figure 3A-B). Metabolomic analysis of the cells and corresponding media were assayed via mass spectrometry (Metabolon, Inc.) and the fold changes of analyzed metabolites were calculated between treatment groups and grouped by metabolic pathway (Figure 4A). Increased glycolysis intermediates, such as glucose-6-phosphate (G6P, p=0.001), fructose-6-phosphate (F6P, p<0.001), pyruvate (p=0.0269) and lactate (p=0.0188), were detected intracellularly in ICOS Strong-activated Th17 cells compared to unstimulated Th17 controls (Figure 4B). Concordantly, Strong ICOS stimulation depleted glucose levels extracellularly and increased lactate secretion compared to Weak ICOS stimulation, suggesting high levels of glycolytic activity (Figure 4C). In contrast, Weak ICOS stimulation enriched isocitrate, phosphoenolpyruvate (PEP), and 3-phosphoglycerate intracellularly (Figure 4B). Furthermore, Weak ICOS stimulation did not induce overt extracellular uptake of either glucose, pyruvate, or glycerate or induce intracellular levels of G6P or F6P suggesting the direction of metabolic flux to favor that of gluconeogenesis instead of glycolysis (Figure 4C).

**Figure 4:**
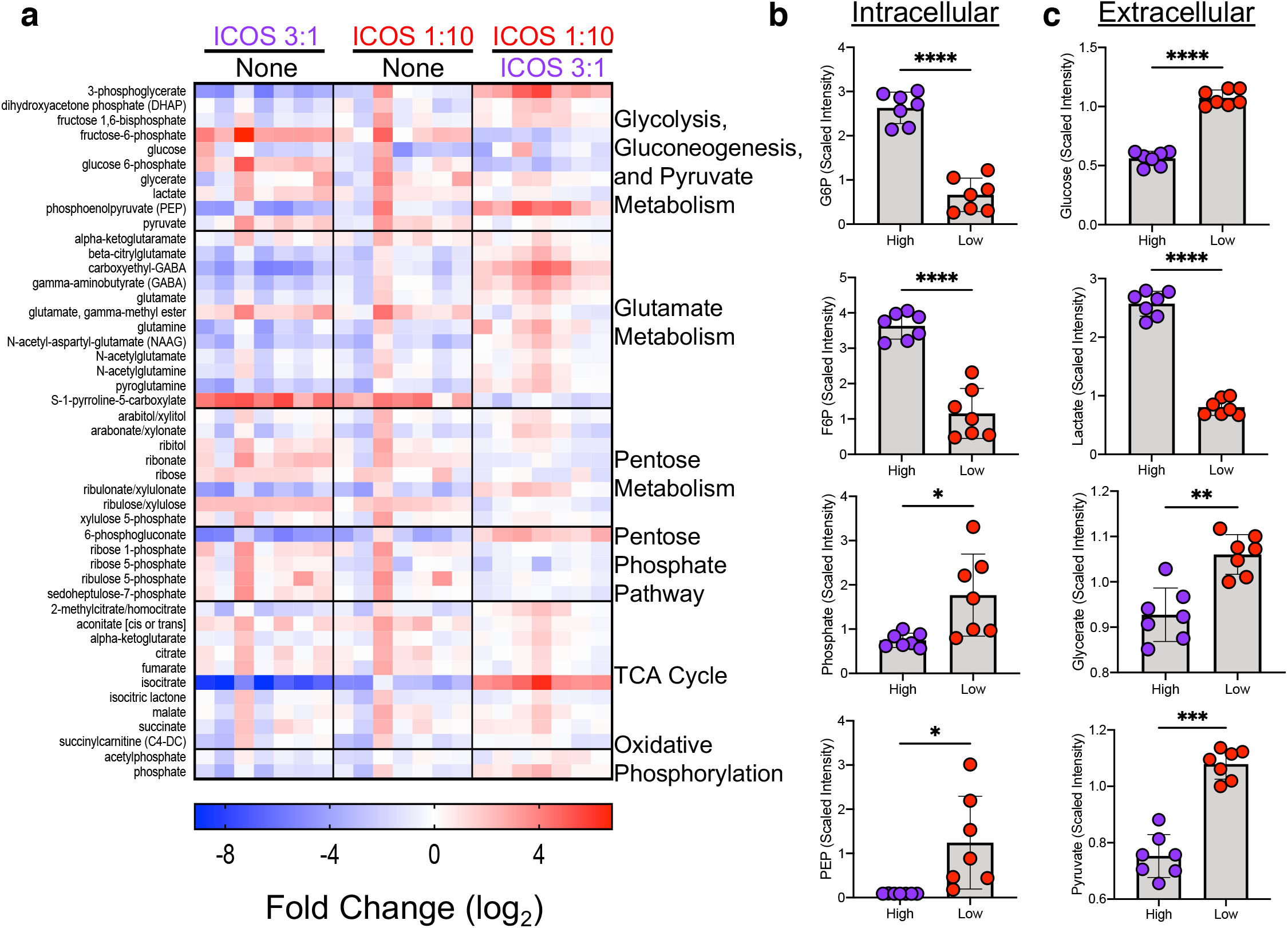
Changes in metabolic states of ICOS Low vs. High activated T cells, compared to no activation, are revealed by metabolomics. Metabolic analysis by mass spectrometry was performed on Th17 cells from healthy donors (N=7) that were activated (or not-”None”) and cultured for 4 days before collection of the cells and their culture media. Resulting values were normalized and fold changes were calculated and log_2_ transformed (f). Normalized values for selected metabolites from the intracellular (g) and extracellular media (h) were averaged. For each assay (a-e), N=3 healthy donors. Statistical analysis was performed using one-way ANOVA with Tukey’s multiple comparisons test (* = P<0.05 ; ** * =P<0.001). Statistical analysis was performed using paired T-tests (* = P<0.05 ; ** =P<0.01; *** =P<0.001).

Comparison of ICOS Weak-versus ICOS Strong-activated Th17 intracellular metabolites revealed further differences in their use of the central carbon pathway, shown by increased abundance of metabolites involved in gluconeogenesis (PEP and 3-phosphoglycerate) as well as the pentose phosphate pathway (6-phosphogluconate), suggestive of active gluconeogenesis and shunting of carbon sources to the penthouse phosphate pathway. An important rate-limiting step of gluconeogenesis is the formation of PEP from oxaloacetate (OAA) catalyzed by the enzyme phosphoenolpyruvate carboxykinase (PCK1) (*30*). Importantly, we found that PEP was 14-fold higher in Weak ICOS-stimulated versus Strong ICOS-stimulated Th17 cells (p=0.0003, Figure 4B). This metabolic profile coupled with the proliferation, differentiation and mitochondrial phenotype led us to hypothesize that Weak ICOS-stimulated cells may be more efficacious when infused into mice bearing a human solid tumor.

### Weak ICOS signal strength augments Th17 cell engraftment and antitumor activity in vivo

Using an established human mesothelioma tumor model, we tested our hypothesis that decreasing the number of activation beads used to generate mesothelin-specific CAR modified Th17 cells *in vitro* would improve their antitumor activity *in vivo*. To do so, bulk human CD4^+^ T cells were programmed toward a Th17 phenotype while being activated with one of 4 treatments, using either αCD3/CD28 or αCD3/ICOS beads at either a 3:1 bead:T cell (“Strong”) or 1:10 bead:T cell (“Weak”) ratio. As shown in the diagram in Figure 5A, NSG mice bearing a human mesothelioma tumor were infused with the MesoCAR Th17 cell products. Human MesoCAR CD8^+^ T cells, expanded with αCD3/CD28 beads at the Strong signal strength (3 beads per T cell), were co-transferred into mice. As anticipated, tumor growth was only slightly and transiently halted in mice treated with CD28 Strong-activated Th17 cells when compared to untreated animals. Tumor growth was controlled for 10 days post-cell infusion in mice treated with CD28-Weak-activated Th17 cells, after which their tumors rapidly expanded (Figure 5B). Mice infused with ICOS co-stimulated Th17 cells mediated superior antitumor responses compared to animals treated with CD28 co-stimulated Th17 cells, consistent with prior reports (*31, 32*). Interestingly, ICOS Strong-activated Th17 cells prevented tumor growth for greater than one month until relapsing, while ICOS Weak-activated Th17 cells imparted durable immunity, marked by sustained control of tumors post-ACT. This striking improvement in Th17 cell response was also reflected by the enhanced survival of ICOS Weak-activated Th17 treated mice (Figure 5C). Donor T cells (%CD45^+^) persisted in the blood of mice two weeks after infusion, revealing that ICOS Weak activation enhanced Th17 engraftment and persistence to the greatest extent compared to other groups (Figure 5D).

**Figure 5.**
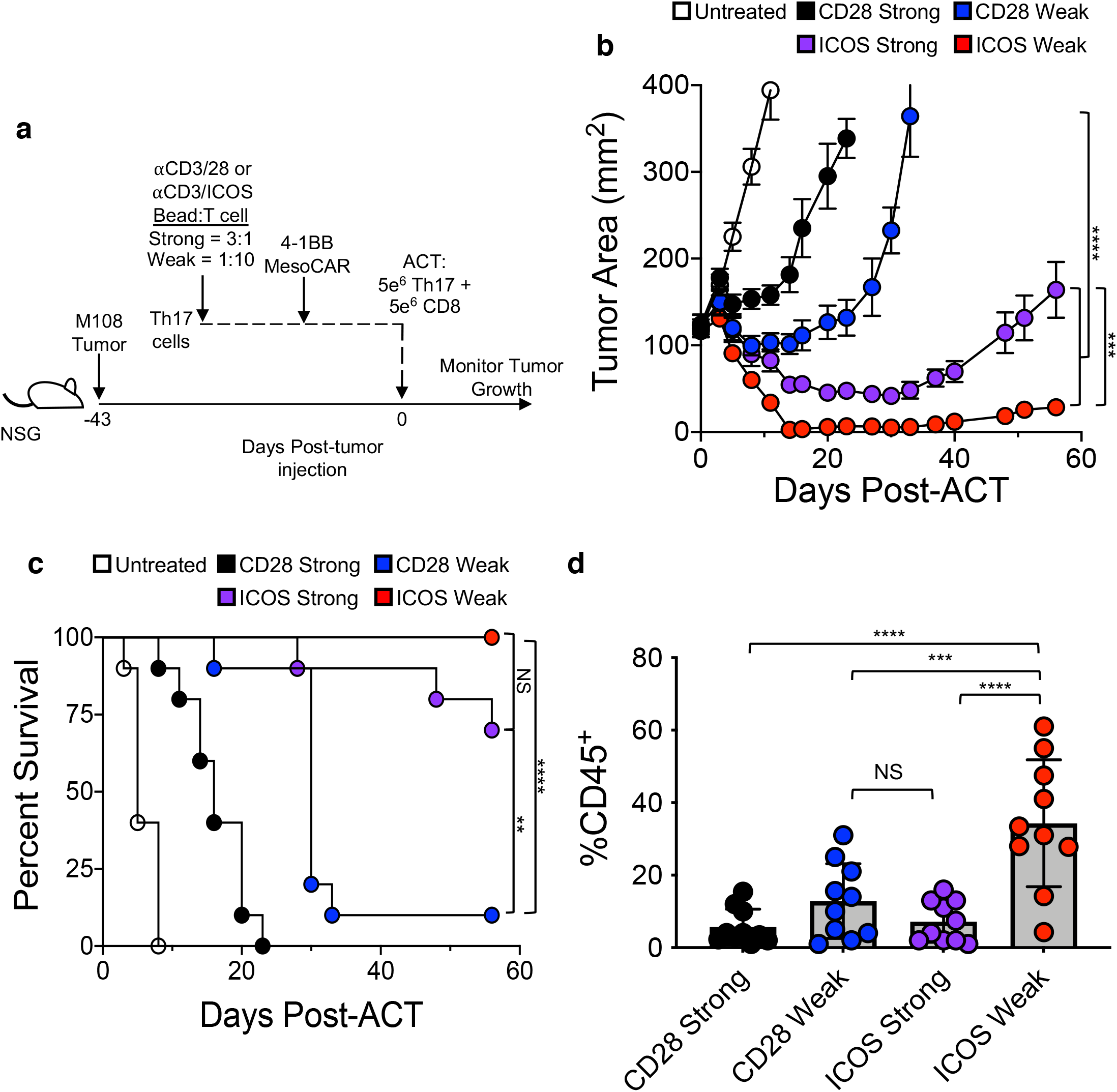
ICOS Low-activated Th17 cells persist *in vivo* and mediate a durable anti-tumor response. a) Experimental schematic. Th17 cells activated with either CD3/28 or CD3/ICOS beads at a 3:1 (“Strong”) or 1:10 (“Weak”) bead:T cell ratio and transduced with the 4-1BB Mesothelin CAR were expanded for 10 days prior to infusion into M108 tumor-bearing NSG mice. 4-1BB Mesothelin CAR CD8 T cells from the same donor were co-infused. Tumor size was measured until mice reached endpoint. Tumor area (mm^2^) (b) and survival to 200mm^2^ tumor area (c) are reported Blood from mice on day 7 post-ACT was processed and analyzed by FACS for presence of human CD45+ cells (d). N=10 mice/group. Statistical analysis was performed using one-way ANOVA with Tukey’s multiple comparisons test and Log-rank Mantel-Cox Test (** =P<0.01; *** =P<0.001; **** =P<0.0001).

## Discussion

T cells can be manipulated in various ways to yield an optimal antitumor cell product for therapy. Th17 cells have particularly demonstrated powerful antitumor prowess (*12-14, 16*), yet the optimal way to expand them has not yet been determined. In the present study, we found that the combination of lower TCR signaling and ICOS costimulation generated a far superior CAR Th17 cell product, with enhanced *in vitro* polyfunctionality, a stem-like memory phenotype and improved antitumor activity. At a mechanistic level, this superior phenotype was mediated by a metabolic switch to oxidative phosphorylation. The concept of decreasing TCR signal strength to improve ACT has been reported previously, whereby decreasing the length of costimulation along with the addition of the cytokine IL-21 expanded stem-like memory T cells (*19*). Similarly, expanding bulk T cells with an equal bead to T cell ratio can promote optimal expansion while maintaining a younger memory phenotype (*20*). Further studies specifically focusing on Th17 cells found that either decreasing the length of time αCD3/28 beads are present in the Th17 culture or by decreasing the number of αCD3/28 beads bolstered their capacity to secrete Th17-associated cytokines, and further elucidated that lowered αCD3/CD28 signal strength promoted Th17 cell responses through the binding of nuclear factor of activated T cells (NFAT) to the proximal region of the IL-17 promotor in low stimulated cells only (*21*). However, modulating the strength of the TCR signal and the co-stimulatory effects of these cells’ ability to impacted immunity against tumors was not tested.

ICOS co-stimulation, compared to CD28 co-stimulatory signaling, enhances Th17 cell generation and antitumor effectiveness (*31, 32*). Herein, we are the first to correlate the concept of reduced TCR signal strength in concert with ICOS signaling to potentiate Th17 cells’ antitumor immunity. CD28 co-stimulatory signaling impairs Th17 function and development, which may impair their therapeutic potential (*33*). Pre-clinical studies of Th17 cells have characterized their advantages, including increased resistance to apoptosis and senescence, and robust antitumor activity, when compared to traditional bulk T cells, Th1 or Th2 cells (*13, 14*). By optimizing their production for use in CAR T cell therapy, we present a way to optimize this therapy for patients with solid tumors.

We found that decreasing the TCR activation signal modulated T cell metabolism in several ways. Metabolically fit Weak αCD3/ICOS-activated cells had lower mitochondrial membrane potential, greater spare respiratory capacity, and decreased glucose uptake, without any loss in cell yield. These data indicate an enhanced usage of oxidative phosphorylation, which has been associated with long-lived memory T cells that have the capacity to produce energy in stressful metabolic conditions, as is often the case in hypoxic tumor microenvironments (*24, 25, 34*). Additionally, OXPHOS has been implicated in bolstering Th17 pathogenicity by supporting mTOR and the BATF transcription factor, which lead to Th17 commitment over Treg development in CD4^+^ T cells (*35*). In contrast, Strong αCD3/28-activated cells sustained high levels of glucose uptake, indicating that they may be more reliant on glucose and glycolysis than Weak activated cells *in vivo*, which could be one detriment to their full functionality in the solid tumor microenvironment.

Weak stimulation of Th17 cells with a 1:10 αCD3/ICOS bead per T cell ratio promoted enhanced levels of metabolites involved in gluconeogenesis (PEP and 3-phosophoglycerate) and the pentose phosphate pathway (6-phosphogluconate). Our metabolite abundance data suggest that compared to ICOS high Th17 cells, which consume extracellular glucose and secrete lactate through glycolysis, ICOS low Th17 likely shunt glutamine or other amino acids towards isocitrate conversion to oxaloacetate and PEP through gluconeogenesis. The lack of abundance of G6P or

F6P and the increase in 3-phosphoglycerate and 6-phosphogluconate suggest the flux of gluconeogenesis towards the pentose phosphate pathway to produce other carbon intermediates and the production of reducing potential, such as NADPH and reduced glutathione to decrease levels of reactive oxygen species. Recent work by has demonstrated that alternative fuels for the central carbon pathway such as inosine can be shunted through the pentose phosphate pathway as a substitute for glucose to improve T cell bioenergetics and tumor immunity (*36*). Thus, Weak ICOS stimulation could alter how T cells use gluconeogenesis for the synthesis of PPP intermediates such as 6-phosphogluconate which is a key intermediate for the generation of both carbon metabolite precursors as well as reducing potential molecules such as NADPH. Our data suggest that alternative usage of the central carbon pathway may be critical in the metabolic rewiring of ICOS Weak-activated Th17 cells and may contribute to their immune potency by providing them with an alternative pathway to supplement glucose in the tumor microenvironment. Our data compliments other reports that PEP sustains NFAT and Ca^2+^ signaling, which results in enhanced effector functions in CD4^+^ T cells (*37*). Complimentary work described upregulated phosphoenolpyruvate carboxykinase (Pck1), the driver of PEP production, in memory CD8^+^ T cells, supporting the pentose phosphate pathway and reduced reactive oxygen species (ROS), supporting memory formation and maintenance (*38*). These data may explain why decreased TCR stimulated cells have increased cytokine function while limiting glucose uptake, as NFAT has also been linked to IL-17 production in low stimulated cells via its binding to the proximal region of the *IL-17* promoter following its translocation to the nucleus (*21*). Furthermore, our Weak ICOS-stimulation data highlights an intriguing alternative rewiring of the central carbon pathway to increase gluconeogenesis and the pentose phosphate pathway to improve the metabolic capacity of Th17 cells. Although the mitochondrial bioenergetic changes observed with Weak ICOS stimulation were subtle compared to Weak CD28 stimulation, this alternative usage of gluconeogenesis seems to be enough to promote their potent antitumor response. Our data supports recent reports showing that supplementation of alternative carbon sources such as inosine, methionine or L-arginine can be sufficient to bolster T cell responses against solid tumors (*36, 39*). We present a novel method to produce human CAR T cells by combining a weak TCR activation signal with ICOS co-stimulation to polarized Th17 cells. This finding has important clinical implications, not only to enhance antitumor response, but to lower the cost of this expensive therapy by drastically reducing the number of antibody-coated beads necessary to generate the cell product with no impact on cell yield. An unknown concern may be that these potent cells may induce associated toxicities such as cytokine release syndrome (CRS) in CAR-treated patients, as has occurred in other potent CAR T cell therapies (*40*), although we did not observe any evidence of this in our mouse models. Based on these studies, we believe this expansion method could improve the cell product and thus the outcome of patients with solid tumor malignancies, leading to long term, durable responses.

## Materials and Methods

### Isolation and Expansion of Human CAR Th17 Cells

Peripheral blood buffy coats were obtained from deidentified healthy donors through the Oklahoma Blood Institute (Oklahoma City, OK). PBMCs were isolated via gradient separation using Lymphocyte Separation Media (Corning), and CD4^+^ and CD8^+^ T cells were further isolated with Dynabeads Human CD4^+^ or CD8^+^ Isolation Kits (ThermoFisher), following manufacturer’s instructions. Cells were activated using Dynabeads M-280 Tosylactivated magnetic beads coated with αCD3 (clone OKT3, Biolegend) and either αCD28 (clone CD28.2, Biolegend) or αICOS (clone ISA-3, eBioscience). CD4^+^ cells were polarized towards a Th17 phenotype at the time of activation with the following conditions: IL-1*β* (10ng/ml), IL-6 (10ng/ml), IL-23 (20ng/ml), α-hIFN-γ (5 μg/ml), and α-hIL-4 (5 μg/ml) and cultured for up to 10 days in culture media (RPMI 1640 with L-glutamine containing 10% heat inactivated FBS, 1% penicillin/streptomycin, 1% non-essential amino acids, 1% sodium pyruvate, 0.1% HEPES, and 0.1% *β*-mercaptoethanol) at 37*°*C, 5% CO_2_. IL-2 (100 IU/ml) and IL-23 (20 ng/ml) were added on day 2 of culture and any time the culture was split thereafter. Cells were de-beaded on day 5 post-activation. CD8^+^ cells were activated with αCD3/28 beads (3 beads per T cell) and cultured in CM with IL-2 (100 IU/ml) for up to 10 days. T Cells were transduced with lentivirus containing the second-generation mesothelin CAR with a 4-1BB costimulatory domain (MesoCAR) vector on day 2 of culture (*41*).

### Flow Cytometry

Cells were collected from culture and washed twice with FACS buffer (PBS containing 2% heat inactivated FBS), and then stained with the appropriate extracellular antibodies (anti-CD45, clone H130, Biolegend; anti-CD3, clone HIT3a, Biolegend; anti-CD4, clone RPA-T4, Biolegend; anti-CD45RA, clone HI100, Biolegend; anti-CD45RO, clone UCHL1, Biolegend; anti-CD62L, clone DREG-56, Biolegend; anti-CCR7, clone G043H7, Biolegend; anti-CD27, clone LG.7F9, eBioscience; anti-CD39, clone A1, Biolegend; anti-Tim3, clone F38-2E2, Biolegend; anti-PD-1, clone EH12.2H7, Biolegend; anti-Glut1, clone 202915, R&D Systems) diluted in FACS buffer for 30 minutes at 4*°*C. For intracellular cytokine staining, cells were restimulated for four hours with PMA/Ionomycin and monensin at 37*°*C, 5% CO_2_ prior to FACS washes and extracellular antibody incubation. After extracellular staining, intracellular staining was performed in fixation and permeabilization buffers (Biolegend) and antibodies at a 1:200 dilution (anti-IFNγ, clone B27, Biolegend; anti-IL-17A, clone N49-653, BD Bioscience; anti-IL-22, clone 2G12A41, Biolegend; anti-IL-2, clone MQ1-17H12). Cells were washed and resuspended in FACS buffer and then analyzed on either an Accuri C6 flow cytometer (BD Biosciences) or a FACSVerse flow cytometer (BD Biosciences). CFSE (ThermoFisher) staining was performed according to manufacturer’s instructions. For mitochondrial stains, cells were resuspended in PBS containing 20nM Mitotracker Deep Red (ThermoFisher) and 250nM tetramethyl rhodamine methyl ester (TMRM, ThermoFisher) for 30 minutes at 37*°*C prior to extracellular staining and flow cytometry. For glucose uptake, cells were resuspended in glucose-free media containing 150 *μ*M 2-(*N*-(7-Nitrobenz-2-oxa-1,3-diazol-4-yl)Amino)-2-Deoxyglucose (2-NBDG Thermofisher) and incubated for 45 minutes at 37*°*C.

### Seahorse

Seahorse extracellular flux analyses were conducted using day 7 cells with the Seahorse XF (Agilient) reagents and XF24 or XF96 instrument, according to manufacturer’s instructions. Briefly, the mitochondrial stress test injected oligomycin (1*μ*M), FCCP (500nM), antimycin (2*μ*M), and rotenone (2*μ*M) while the instrument measures the oxygen consumption rate (OCR). Percent spare respiratory capacity was calculated using the manufacturer’s spreadsheet macros.

### Mice and Tumor Models

NOD/*scid* gamma chain knockout (NSG) mice were obtained from Jackson Laboratories and maintained in the animal facilities of the University of Pennsylvania School of Medicine, in accordance with IACUC regulations. The human mesothelioma cell line, M108, was cultured in E media (RPMI 1640 containing L-glutamine with 0.5% Human serum, 1X ITES, 10mM HEPES, 0.5mM sodium pyruvate, 0.1mM non-essential amino acids, 1X Pen/Strep, 1ng/mL recombinant human epithelial growth factor, 18ng/mL hydrocortisone, and 0.1 nM triiodothyronine) for up to 2 passages. 5 × 10^6^ M108 cells in a 1:1 mixture of PBS and Matrigel (Corning) were subcutaneously injected into NSG mice. Tumors were allowed to establish for 43 days, and then Th17 CAR T cells combined with CD8^+^ CAR T cells were adoptively transferred into tumor-bearing mice via tail vein injection. Prior to therapy, mice were assigned randomly to groups based on tumor size, and L x W measurements via calipers were collected by personnel blinded to treatment group. Experimental endpoints were determined prior to study execution. Tumor control animal experiments were conducted over ∼60-70 days. Animals were only excluded if tumors were very small or immeasurable prior to therapy initiation. Outliers were reported. Tumor end point was established at 400mm^2^.

### Metabolomic Analysis

Th17 cells from 7 normal healthy donors were activated with either Strong ICOS bead stimulation (3 beads per T cell), Weak ICOS bead stimulation (1 bead per 10 T cells), or were left inactivated (no beads). Cells were cultured for 4 days and then harvested. The cells were removed from their cell media by centrifugation and then frozen at -80*°*C along with the separated cell media. Metabolon, INC (Morrisville, NC) performed metabolomic analysis using mass spectrometry and bioinformatic analysis.

### Statistical Analysis

For *in vitro* experiments, a sample size of ≥3 was chosen. For *in vivo* experiments, a sample size of ≥5 was chosen. Statistical analysis was performed using GraphPad Prism Software v9 (GraphPad Software, LLC). Experiments comparing two groups were analyzed using a Student’s *t*-test. For experiments comparing three or more groups, a one-way analysis of variance (ANOVA) was performed with a post-comparison of each group using the Tukey’s Multiple Comparisons test. Graphs utilizing error bars display the mean as the center value and the error bars indicate SEM. For tumor curve experiments, a one-way ANOVA with Tukey’s Multiple Comparisons test was performed at the final dates where all mice from the compared groups remain alive. Survival curve analysis was performed using a Log-Rank Mantel-Cox test. For the metabolomics study, fold change from the ‘No Stimulation’ group values were calculated and log_2_ transformed and reported in the heat map.

## Contributors

MMW: Conceptualization, data curation, formal analysis, investigation, methodology, visualization, writing—original draft, writing—review and editing. LWH: Conceptualization, data curation, investigation, methodology, writing—review and editing. MHN: Conceptualization, data curation, investigation, methodology, supervision, formal analysis, writing—review and editing. LRN: Conceptualization, data curation, investigation, methodology, writing—review and editing. ARM: Data curation, investigation, methodology, writing—review and editing. GORR: Data curation, investigation, validation, writing—review and editing. ASS: Data curation, investigation, writing—review and editing. AMRR: Data curation, investigation, writing—review and editing. HMK: Data curation, investigation, writing—review and editing. JLR: Resources, supervision, writing—review and editing. GRL: Data curation, formal analysis, supervision, writing—review and editing. CMP: Conceptualization, data curation, formal analysis, funding acquisition, investigation, project administration, resources, supervision, visualization, writing—original draft, writing—review and editing, guarantor.

## Acknowledgements

Supported in part by the Flow Cytometry and Cell Sorting Shared Resource, Hollings Cancer Center, Medical University of South Carolina (P30 CA138313).

